# Giant Viruses and their mobile genetic elements: the molecular symbiotic hypothesis

**DOI:** 10.1101/299784

**Authors:** Jonathan Filée

## Abstract

Among the virus world, Giant viruses (GVs) compose one of the most successful eukaryovirus families. In contrast with other eukaryoviruses, GV genomes encode a wide array of mobile genetic elements (MGEs) that encompass diverse, mostly prokaryotic-like, transposable element families, introns, inteins, restriction-modification systems and enigmatic classes of mobile elements having little similarities with known families. Interestingly, several of these MGEs may be beneficial to the GVs, fulfilling two kinds of functions: 1) degrading host or competing virus/ virophages DNA and 2) promoting viral genome integration, dissemination and excision into the host genomes.

By providing fitness advantages to the virus in which they reside, these MGES compose a kind of molecular symbiotic association in which both partners should be regarded as grantees. Thus, protective effects provided by some of these MGEs may have generated an arms race between competing GVs in order to encode the most diverse arsenal of anti-viral weapons, explaining the unusual abundance of MGEs in GV genomes by a kind of ratchet effect.

**Highlight:**

- Giant Virus (GV) genomes are loaded with diverse classes of mobile genetic elements (MGEs)
- MGEs cooperate with GV genes in order to fulfill viral functions.
- Site-specific endonucleases encoded by MGEs are used as anti-host or anti-competing viral compounds
- Integrase/transposase genes derived from MGEs have been recruited to generate integrative proviral forms.
- MGEs and GVs may thus compose a mutualistic symbiosis

## Introduction

Giant Viruses (GVs) compose a remarkable group of eukaryoviruses: prevalent in many environments from the deep ocean to the human gut, they infect a wide spectrum of hosts ranging from tiny protists to vertebrates. They possess highly variable genomes in term of size and content (from 100 kb to 2,5Mb), play important ecological roles in controlling hosts proliferation and are even suspected to be human pathogens [1–6]. Thus, this peculiar group of virus has been successful in nature and it is important to understand the reasons of this success.

Compared to other eukaryoviruses, GV genomes display two salient features. The first one is the abundance of genes horizontally acquired from cellular organisms. In average, 10% of the GV genes have been acquired by this process, mainly from prokaryotic sources and occasionally from their hosts [7–11]. The second special feature is the level of gene duplication. For instance, the largest genome of GVs has more than 50% of its sequence occupied by genes belonging to duplicated families [16].Thus, there is a positive correlation between the GVs genome size and the number of repeated genes [12–15]. Experimental data indicate also that there is a dynamic process of gene duplication and gene loss in response to host adaption [17,18]. All together, importance of horizontal gene transfers and lineage-specific gene duplication have led to the emergence of the «genomic accordion» hypothesis [12,17,19,20] which proposes that GVs adapt to new niches, hosts or changing environments by series of genome contraction and expansion using a combination of gene accretion, duplication and deletion. Whereas this model need to be supported by additional experimental and comparative genomics evidences, the alternative scenario implying a common ancestry with cells has lost some audience. Indeed, the existence of a «fourth kingdom of life» lack reliable phylogenetic signals supporting the kinship of GVs with cells [21] in addition to the absence of genomic proof of a general tendency of GVs genomes to decrease in size and in diversity with time [7,19,22]. At the opposite, the « genome accordion » scenario fit perfectly well with a model of genome size evolution observed in birds and mammals under the cross effect of expansion and loss of mobile genetic elements (MGEs) [23].

MGEs form a disparate group of selfish genes that tend to act as molecular parasites into the genomes in which they reside. They are abundant in cellular genomes [24] but rare in the virus world and even almost absent in eukaryovirus [25]. Strikingly, GVs are the exception to this rule. MGEs are diverse and abundant in GV genomes and GVs have apparently cope with this load for a long period of time [9,15,19]. In this paper, I collect multiple evidences that several MGEs found in eukaryoviruses have been recruited to perform viral functions. I show that the MGE/GV association has evolved towards cooperation rather than competition, allowing the MGEs to reside in a stable form in the GV genomes. This association, by providing selective advantages to the GVs also promote the MGEs dissemination. Thus, I propose the hypothesis that the unusual abundance of MGEs in GV genomes should be explained by the existence of a kind of molecular symbiotic relationship in which both partners benefit from the presence of each other’s.

## Distribution and origin of MGEs in eukaryoviruses

Phylogenies of hallmark genes and phyletic distributions of the genomic repertoires indicated that at least nine GV lineages exist (Figure 1) [26]. Most of these viruses harbor a large diversity of MGEs (Figure 1, Table 1). The majority of them are transposable elements, including diverse Insertion Sequence (IS) families and Eukaryotic-like DNA transposon families as Mariner, PiggyBac or Polinton/Virophage. The last family is of special interest as it clusters a huge diversity of elements including satellite viruses of the GVs (Virophages), composite transposons (Polintons) and their highly-reduced derivatives (Transpovirons) [27–29]. It shall be noted that the capsid protein of the Virophages is related to that of the GVs, suggesting strongly that these elements share a common ancestor [27].

**Table 1:**
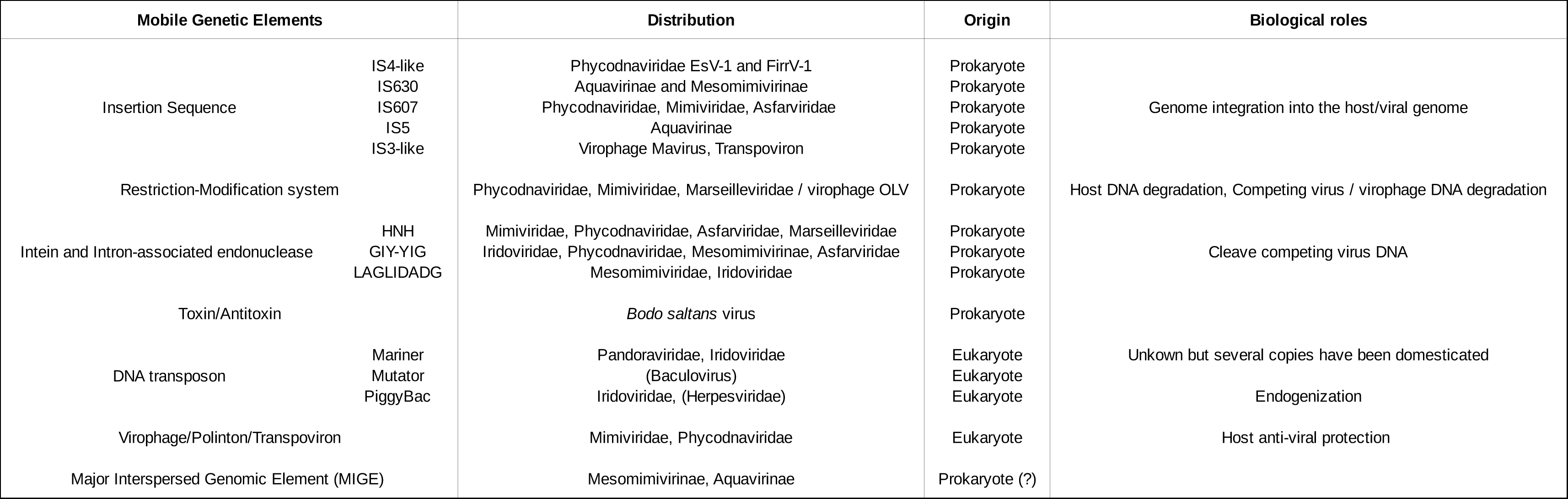
Distribution, origin and possible biological roles of mobile genetic elements found in virus genomes.

**Figure 1:**
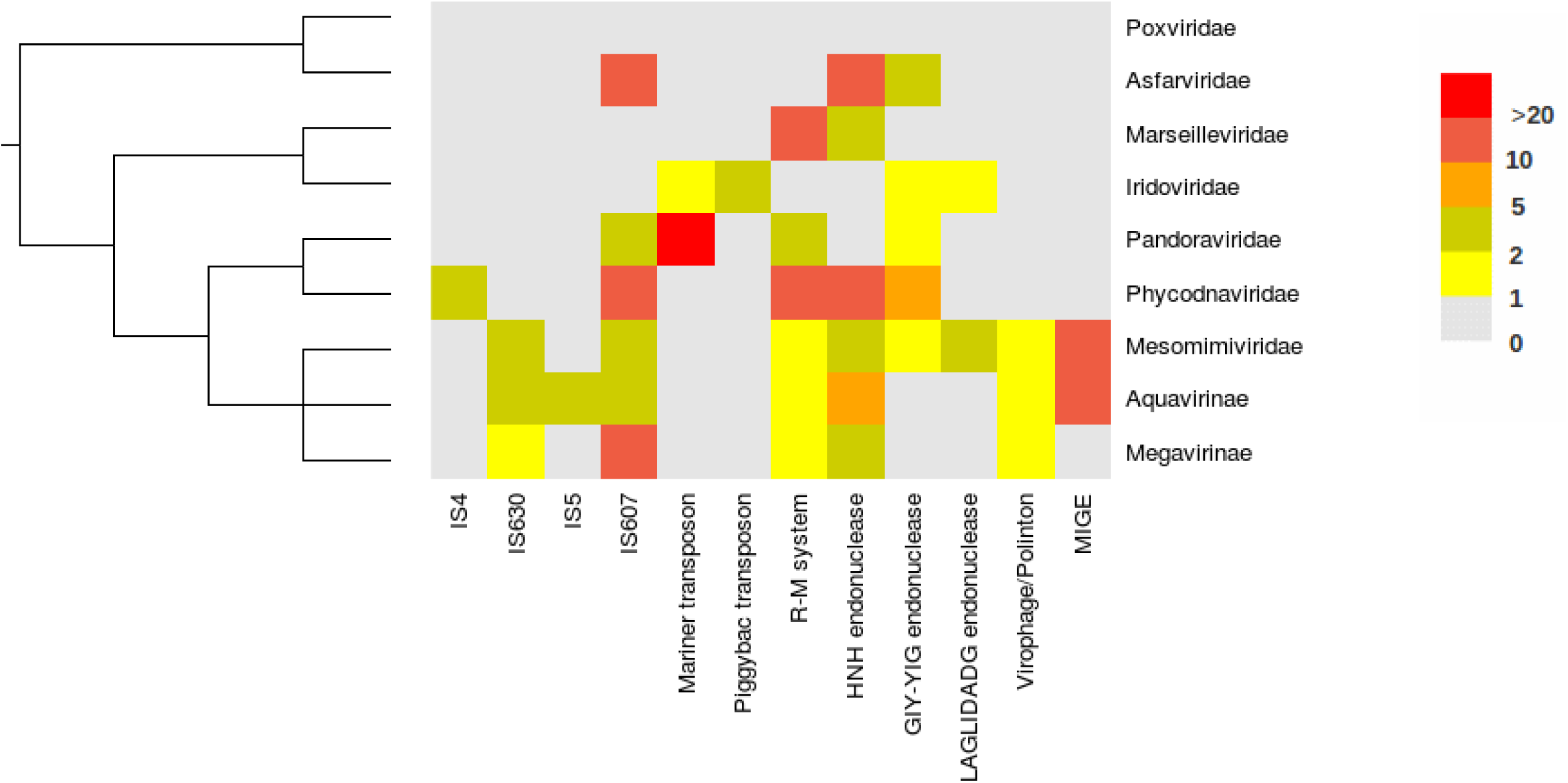
Diversity of mobile genetic elements found in Giant Virus genomes. The heat-map represents the maximum number of copies of each family of mobile elementsidentified in each viral lineages. The phylogeny is a consensual tree obtained with conserved viral core genes.

In addition to transposable elements, there are also two kind of Restriction/Modification systems (R-M systems) composed by a gene tandem encoding a restriction endonuclease and a methyltransferase. The endonuclease recognizes and cleaves at specific sites foreign DNA whereas the methyltransferase protects self DNA by transferring methyl groups at the cleavage sites. These systems considered as primitive immune systems in prokaryotes are known to behave as selfish mobile elements. In the case of a system loss, the endonuclease tends to persist in the cytoplasm for a longer period of time compared to the methyltransferase, leading to the restriction of GV and host genomes [30]. R-M systems are not only stably maintained in genomes as addictive selfish modules but also tend to propagate efficiently by lateral gene transfers [31]. Recently, a new class of selfish mobile element has been evidenced in the *Bodo saltans* virus (Aquavirinae) [26]. The toxin-antitoxin systems (T-A) behave in a similar way as R-M systems and also tend to become addictive modules. In addition, most of GV genomes encode for numerous mobile homing endonucleases frequently associated with self-splicing introns and inteins (also known as protein introns) [9,32,33]. Homing endonucleases are mobile and selfish genes that are known to be invasive, sometimes promoting the «homing» of their cognate introns or inteins into empty sites by the mean of homologous recombination [34].

Finally, more enigmatic MGEs have been evidenced in Mimiviridae: the Major Interspersed Genomic Elements (MIGEs) [32,35]. These elements are composed of a single ORF encoding a protein with a zinc-finger motif preceded by a short piece of non-coding DNA. MIGEs are present in multiple copies (up to 20) and the ORFs have some homologs in a limited number of bacterial genomes. Their mode of dissemination is still unknown but they have successfully invaded at least 6 different GV genomes belonging to the Aquavirinae and Mesomimivirinae.

The patchy distribution of most of these MGEs (Figure 1) suggest complex evolutionary histories with multiples events of loss and gain. Most of these MGEs have only prokaryotic homologs suggesting that they have been acquired by lateral transfers from Bacteria or Archaea (Table 1). However, phylogeny of the IS families indicates that GV sequences form monophyletic clusters, suggesting a single event of acquisition for each family [8]. Finally, the genomic comparison of closely related GVs shows the existence of frequent movements and dissemination of these MGEs in addition to vertical and stable transmission for some others ones [19].

Such abundance, diversity and apparent long-standing association of MGEs in GV genomes is a very unusual feature in eukaryoviruses [25]. A possible explanation of this pattern is the existence of reciprocal benefits underlying the association of MGEs with GVs, as suggested by the literature which reveals two kinds of mutualistic association between MGEs and GVs that will be described in the next sections.

## Association of MGEs and viruses to degrade host or competing virus genomic DNA

The first direct evidence of cooperation of MGEs and GVs has came from the unusual presence of numerous Restriction-Modification systems in *Chlorella* Phycodnaviruses (Figure 1). These prokaryotic MGEs are known to cooperate with diverse bacterial species in order to degrade invading phage DNA [30]. In Phycodnaviruses, it was shown that the endonuclease of the R-M system is used to degrade immediately after infection the host genomic DNA whereas the methyl-transferase protects the viral DNA [36]. By procuring an abundant source of deoxynucleotides that can be reincorporated into the viral DNA, R-M systems and GVs compose a dual association with an increased global fitness. Do the R-M systems also protect against competing viruses by degradation of their unprotected genomes? So far, this is not evidenced, however in the case of simultaneous infections by two different GVs, a mutual exclusion has been reported in *Chlorella* Phycodnaviruses [37]. Moreover, the Organic Lake Phycodnavirus (OLPV) encodes a R-M system for which the cognate methyl-transferase, but not the endonuclease, is also encoded by the virophages OLV that are known to infect the OLPVs [38]. Taken together, these data support the view that R-M systems might also be used as anti-viral/anti-virophage agents, leading to an arms race between different co-infecting GVs and/or between GVs and virophages in order to encode the most diverse set of R-M systems.

GVs also encode numerous introns and inteins that are often associated with a site-specific homing endonuclease. It was demonstrated in diverse bacteriophage groups that these elements are implicated in exclusion processes during competitive infection by two or more viruses [39–41]. By disrupting the enzyme recognition site, the intron prevents self-cleavage by the endonuclease. However, during a mixed infection by two related viruses that differs by the presence or the absence of an endonuclease associated with introns inserted in the same gene, the intron-free genes are specifically cleaved, reducing the fitness of the corresponding virus and favoring the viral genome in which the intron is present [26]. Supporting this view, it’s noteworthy that inteins and introns are most often inserted in conserved GV «core» genes that are known to be essential for the achievement of the virus cycle such as DNA polymerase, Ribonucleotide reductase, RNA polymerase or Capsid genes [33,42–44]. Additionally, it was evidenced that the well-known strong site-specificity of the homing-endonuclease is relaxed in GV genomes allowing the inteins to be inserted at multiple sites, in different genes, widening their possible anti-viral effects [32]. These data support the view that the multiplication of introns or inteins in GV genomes are not simply the result of proliferation of selfish genetics elements. They are more likely the consequence of an arm race between competing viruses that have co-opted these genomic parasites as anti-viral weapons.

In this framework, the endonuclease encoded by R-M systems or associated with introns or inteins should be viewed as the anti-viral, anti-virophage or anti-host poison whereas the methyltransferase or the intron or the intein themselves were the antidotes. Incidentally, these MGEs have become addictive modules, applying strong selective pressures favoring their fixation in the viral population and boosting their horizontal dissemination. For these reasons, the association of R-M systems, introns and inteins with the GVs would compose a kind of molecular symbiosis by providing benefits to each partners.

## MGE and virus cooperate in order to promote provirus integration

Viral genome integration into the host genome is a frequent phenomenon in viruses. Indeed, for some viruses, the viral DNA is integrated into the host genome as a transient step of their lifecycle (for example retroviruses) whereas other ones have evolved stable «proviral» versions into host genomes that are transmitted vertically from the parents to the progeny (see for review REF [45]). This process is called endogenisation when the virus has ultimately loss is capability to generate infectious viruses. Interestingly, MGEs seem to have promoted both processes during eukaryovirus evolution.

Some GVs belonging to the Phycodnarividae family integrated their genomes into their algal host genomes as a biological step of their life cycle [46,47]. The genome sequence of two GVs that generated provirus has revealed the presence of a XerC-like integrase and DDE transposases in both genomes. These enzymes are suspected to catalyze the integration and the excision of the viral DNA into the host genomes [48,49]. Moreover, multiple pro-viral copies of different sizes and inserted at specific GC-CG insertion sites co-exist in the host genomes, suggesting that viral genome integrations are not the result of accidents [47,50]. In this case, the domestication of the integrases/transposases has allowed the viruses to generate a «sleeping» version that is adapted to the peculiar lifestyle of their algal hosts, as Ectocarpale alga have complex life cycles with vegetative and reproductive stages. Viral particles are only released in the environment during the reproductive steps when spores or gametes are generated [51]. If the latency process observed in Ectocarpale viruses is unusual in GVs, the extant of the phenomena might be underestimated as a nearly complete mimiviridae-like virus genome has been found integrated into the *Hydra* genome [52] and multiple GV genomes fragments have been reported in various cellular genomes [52,53].

The endogenization process should be viewed as the ultimate stage of the latency. The virus has lost its infectious property and behave as selfish DNA as observed for transposons. However, in order to efficiently resides and propagates into the host genome, the virus needs enzymes to catalyze the excision/integration step. Some Herpesviruses, which do not belong to the GV group despite their large genome size (up to 300 kb) have resolved the issue by co-opting a transposase derived from a PiggyBac transposable elements [54,55]. The resulting elements called ‘Teratorn’ lead to large endogenous viral genomes (up to 200kb). They are sometimes surrounded by typical inverted repeats found in PiggyBac transposon that are probably recognized by the transposase to mediate the dissemination of the elements. Teratorn elements have successfully invaded a wide variety of fish genomes in which they are present in high copy number (>20 copies) [56]. Thus, Teratorns provide an additional case of cooperation between a virus and a transposon, in order to successfully invade host genomes by generating endogenous versions of the virus.

A very similar scheme is also observable in Virophages. As previously stated, Virophages are distantly related GV cousins that have in turn became parasites of their relatives. The relationships of Virophages with Polinton and Transpovirons (their miniaturized derivatives) are very complex and look like a network of gene gains and losses [57]. Virophages have the capabilities to insert their genomes both into the GV [29] and the cellular host genome [3,58]. Provirophage genes integrated into the cellular genomes are specifically activated during GV infection conferring an anti-viral protection to the host cells by inhibiting the GV replication. Conversely, it’s not known if the provirophage integration into the GV genomes increases the fitness of the association by conferring a similar protection against competing virophages for instance. The cooperation between cells and virophages is promoted by the integration and the subsequent dissemination of provirophages into the cell genomes. This process has been achieved by co-opting diverse integrase and transposase genes derived from MGEs. Sputnik-like virophages encode a phage-type, tyrosine recombinase-like integrase and a IS3-like transposase, whereas most Polinton transposons and Mavirus virophages encode a DDE integrase [27,58–60]. Transpoviron genomes do not encode any integrase but an IS3-like transposase gene is present. This transposase has some similarities with the Sputnik IS3 transposases [29]. The mutualistic association between virophage-like elements and MGE genes domesticated to mediate provirophage integration/dissemination provide an additional remarkable example of tripartite cooperation between the host cell, the virophage and the domesticated integrase/transposase genes in order to suppress viral infection, increasing the fitness of the symbiotic system.

## Conclusion

The abundance and the diversity of MGEs is one of the most salient features of the GVs genomes. The successful invasion of these MGEs should be explained by the existence of a kind of molecular symbiosis: while the GV provides a niche in which the MGEs are stably maintained and vertically transmitted in the viral progeny, MGE genes has been used to fulfill viral functions as defense against competing viruses, degradation of the host DNA or the generation of provirus able to insert and reside into the host genomes. By providing competitive advantages to the GV, the MGEs inserted into the GV genomes also increase their own fitness. However, R-M systems and intron/intein associated with homing endonucleases may also became addictive as the loss of the methyltransferase genes or the introns/inteins would no longer protect the viral genome against the deleterious effect of the corresponding site-specific restriction enzymes. Competition between different viruses infecting the same host may also lead to an arm race in which each virus have deployed an arsenal of anti-viral compounds. As new MGE families are frequently discovered in GVs genomes [35,61] new kinds of association may be discovered in the next future providing further supports to the existence of molecular symbiosis between these genetic entities.

## Acknowledgments

Jean-Michel Rossignol for providing useful comments to the manuscript.

